# *cdon* and *boc* affect trunk neural crest cell migration non-cell autonomously through a reduction of hedgehog signaling in zebrafish slow-twitch muscle

**DOI:** 10.1101/2022.01.10.475733

**Authors:** Ezra Lencer, Rytis Prekeris, Kristin Artinger

## Abstract

The transmembrane proteins *cdon* and *boc* are implicated in regulating hedgehog signaling during vertebrate development. Recent work showing roles for these genes in axon guidance and neural crest cell migration suggest that *cdon*/*boc* may play additional functions in regulating directed cell movements. We use novel and existing mutants to investigate a role for *cdon* and *boc* in zebrafish neural crest cell migration. We find that single mutant embryos exhibit normal neural crest phenotypes, but that neural crest migration is strikingly disrupted in double *cdon*;*boc* mutant embryos. We further show that this migration phenotype is associated with defects to the differentiation of slow-twitch muscle cells, and the loss of a Col1a1a containing extracellular matrix, suggesting that neural crest defects are a secondary consequence to defects in mesoderm development. Combined, our data add to a growing literature showing that *cdon* and *boc* act synergistically to promote hedgehog signaling during vertebrate development, and provide a foundation for using zebrafish to study the function of hedgehog receptor paralogs.

## Introduction

During vertebrate development, neural crest cells (NCCs) migrate ventrally from the dorsal neural tube along conserved pathways in the embryo to contribute to numerous traits such as the craniofacial skeleton, peripheral nervous system, and pigment (Theveneau and Mayor, 2012). Defects in NCC migration result in multiple human hereditary disorders (Tobin et al., 2008; Vega-Lopez et al., 2018), and modifications to NCC migration is associated with the evolution of morphological diversity among some vertebrate taxa (Powder et al., 2014). Further, diseases, such as metastatic melanoma, have been shown to progress through a reactivation of a NCC-like migratory genetic program (Diener and Sommer, 2021; Kaufman et al., 2016). Thus, understanding the genetic regulation of NCC migration is fundamental to understanding the etiology to multiple human diseases as well as a major developmental event that occurs in every vertebrate embryo.

NCC migration is highly coordinated along conserved routes in the embryo (Mayor and Etienne-Manneville, 2016), and is regulated by both NCC intrinsic factors, such as genes regulating cell adhesion and actin cytoskeleton (Clay and Halloran, 2011, 2013, 2014; Williams et al., 2018), as well as cell extrinsic factors such as proteins in the extracellular matrix (Perris and Perissinott, 2000; Szabo et al., 2016), tissue stiffness (Shellard and Mayor, 2021), and chemical gradients (Barriga et al., 2013; Olesnicky Killian et al., 2009; Shellard et al., 2018) that funnel NCCs along the migratory route. Mediating cell intrinsic and extrinsic regulation of NCC migration are cell signaling interactions (Carmona-Fontaine et al., 2008; Mayor and Etienne-Manneville, 2016; Scarpa et al., 2015; Szabo et al., 2016; Theveneau and Mayor, 2012). These include the chemical gradients that attract and inhibit migrating NCCs, as well as adhesion molecules that adhere NCCs to each other and mediate cell-cell interactions such as contact inhibition of locomotion (Carmona-Fontaine et al., 2008; Mayor and Etienne-Manneville, 2016; Scarpa et al., 2015; Szabo et al., 2016; Theveneau and Mayor, 2012; Vega-Lopez et al., 2017).

Work has implicated a role for the immunoglobin superfamily member protein *cell adhesion associated, oncogene regulated* (*cdon*) as a cell surface protein regulating trunk NCC (tNCC) migration in zebrafish (Powell et al., 2015). Morpholino knockdown of *cdon* results in arrested (tNCC) migration, with tNCCs mislocalizing N-cadherin (Cdh2). *cdon* and its paralog *boc, brother of cell adhesion associated, oncogene regulated* (*boc*), are vertebrate members of a larger gene family with identifiable orthologs in most metazoan taxa including the *Drosophila* paralogs *ihog* and *boi*. Recent work has established a role for these proteins in activating hedgehog signaling. *cdon* and *boc* act as co-receptors binding to hedgehog ligand and patched receptor to promote smoothened activity. Other work suggests that *cdon*/*boc* may also interact with Wnt and Nodal signaling pathways (Hong et al., 2020; Jeong et al., 2017). *cdon* has been shown to form a complex with N-cadherin/Cdc42/Bnip-2 in a hedgehog independent manner to regulate actin cytoskeletal dynamics during myogenic differentiation in cell culture (Kang et al., 2008; Lu and Krauss, 2010). *boc* has been shown to play a role in axon guidance neuron growth (Connor et al., 2005; Okada et al., 2006). Both proteins localize to the ends of long membrane extensions (e.g. cytonemes), suggesting that these receptors help cells sense the external environment and communicate over long distances (Bilioni et al., 2013; Ferent et al., 2019; Hall et al., 2021). We hypothesize that these transmembrane proteins are reiteratively used as cell surface proteins to integrate multiple signals during processes such as cell migration and axon guidance. While a role for *cdon* in tNCC migration has been suggested (Powell et al., 2015), the mechanism by which *cdon*/*boc* regulates NCC movements is not fully understood.

In zebrafish, hedgehog signaling is thought to affect tNCC migration in a non-cell autonomous manner by regulating the differentiation of adaxial mesoderm (Honjo and Eisen, 2005). Adaxial cells form at the border of the notochord during gastrulation and differentiate into slow-twitch muscle cells that migrate laterally through the somites (Elworthy et al., 2008; Wolff et al., 2003). A non-migratory population of adaxial cells, called muscle pioneers, remain at the body midline. In hedgehog mutant embryos, adaxial cells fail to differentiate into slow-twitch muscle/muscle pioneer derivatives (Elworthy et al., 2008; Roy et al., 2001; Yin et al., 2018). Zebrafish embryos with slow-twitch muscle defects also exhibit tNCC migration defects. Though the mechanisms of how these two tissues interact is not fully understood, it is suggested to depend on extracellular matrix deposition/modification (Banerjee et al., 2011; Banerjee et al., 2013; Guillon et al., 2016; Honjo and Eisen, 2005). Whether *cdon*/*boc* affect slow-twitch muscle differentiation is unknown (though see (Bergeron et al., 2011)), and it is possible that *cdon*/*boc* affect NCC development through autonomous mechanism (Powell et al., 2015).

We use novel CRISPR/Cas9 generated mutations and existing mutants to investigate the role of *cdon* and *boc* in tNCC migration and slow-twitch muscle differentiation. We show that *cdon* and *boc* are expressed by tNCCs and adaxial mesoderm. Loss of either *cdon* or *boc* individually has no observable effect on tNCC migration and minimal effects on slow-twitch muscle differentiation. However, embryos that are double mutant for *cdon* and *boc* exhibit both tNCC migratory defects and loss of slow-twitch muscle. We show that tNCC mobility is not impaired by loss of *cdon*/*boc* suggesting that the tNCC migration defect is a non-cell autonomous consequence of loss of the migratory path. We further provide data suggesting that *cdon*/*boc* are modifying hedgehog signal reception during mesoderm differentiation. These data contribute to a growing literature on the critical role of *cdon*/*boc* in modulating hedgehog signaling across multiple tissues during embryogenesis.

## Results

### cdon/boc paralogs exhibit shared sequence conservation in extracellular domains

*cdon* and *boc*, share a similar genomic domain architecture that includes 4 immunoglobin (IG) domains, 3 fibronectin type 3 repeat (FNIII) domains, and a cytosolic C-terminal tail (**Figure 1A**). A fifth IG domain is variable present in *cdon* orthologs of tetrapod taxa (grey in **Figure 1A**). The FNIII domains have been implicated in binding to hedgehog ligand and patched receptor (Izzi et al., 2011; Song et al., 2015), and evidence suggests that the c-terminal region is needed for protein localization in the cell (Okumura et al., 2021).

**Figure 1.**
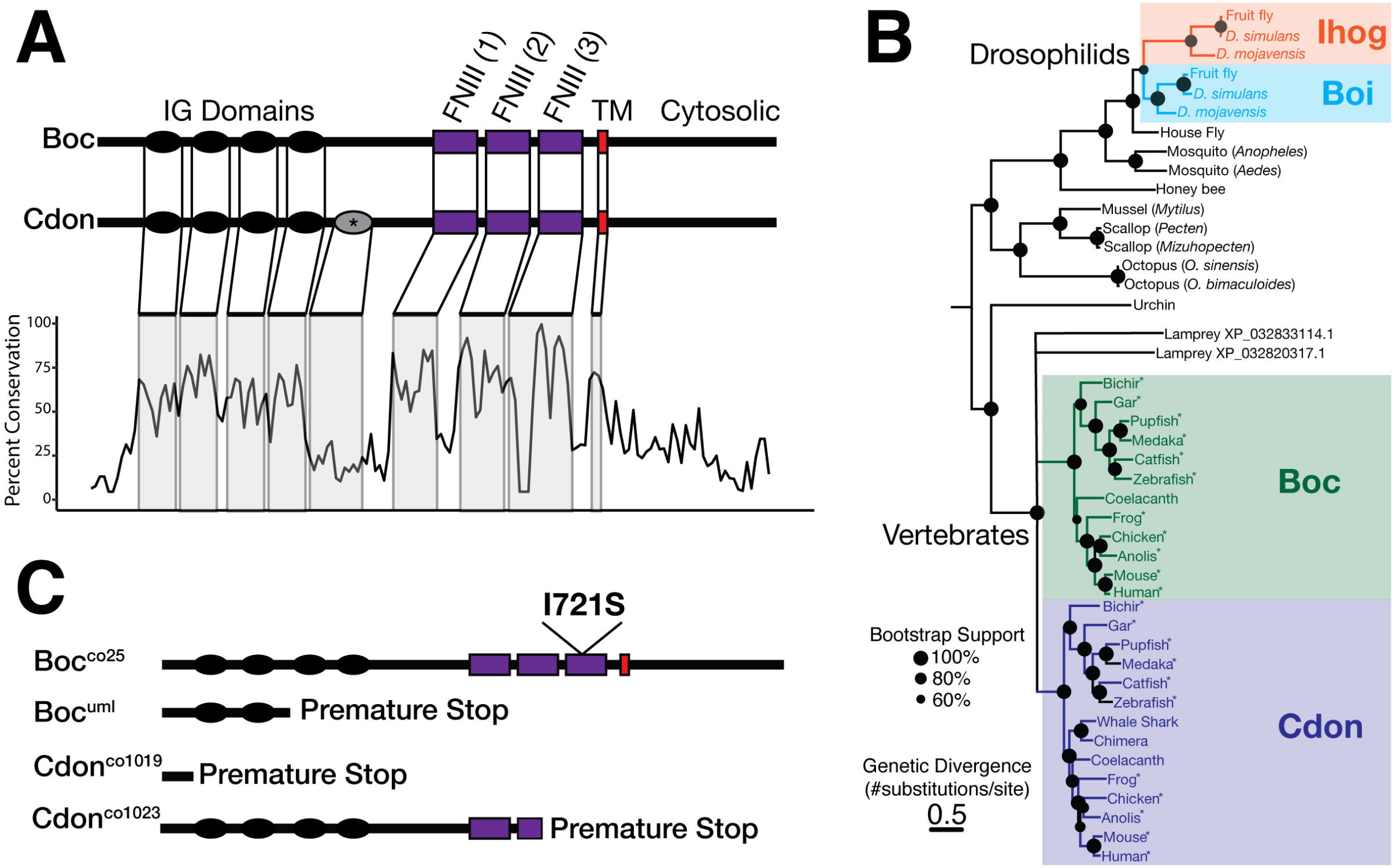
*cdon* and *boc* are transmembrane receptors and share conserved domain structure. **(A)** *cdon*/*boc* share a domain structure of 4-5 immunoglobin (IG) domains, 3 fibronectin (FNIII) domains, a transmembrane (TM) region, and a cytosolic C-terminal tail. A fifth *cdon* IG domain (*) is found in some tetrapod taxa. Percent conservation (20 AA sliding window) is shown for an alignment of *cdon* and *boc* amino acid sequences from 11 tetrapod and ray-finned fish taxa. Note that the extracellular domains exhibit high sequence conservation, while the cytosolic region is less conserved. **(B)** Maximum likelihood tree (phyml) for curated *cdon*/*boc* ortholog amino acid sequences from multiple bilaterian taxa. Note independent duplications for Ihog/Boi in drosophilids and Cdon/Boc in vertebrates. Node support was calculated based on 1000 bootstraps, and nodes with less than 60% bootstrap support were collapsed. Asterisks mark taxa used to generate conservation in panel A. **(C)** Schematic of *cdon* and *boc* mutant alleles characterized in the current study.

As functional conservation is likely to be reflected by sequence conservation across evolutionary time, we investigated the molecular evolution of *cdon*/*boc* orthologs across vertebrate taxa. An alignment of *cdon* and *boc* amino acid sequences from 11 select jawed vertebrate taxa shows that extracellular domains exhibit high conservation across taxa and paralogs, suggesting that these extracellular domains are critical for gene function (**Figure 1A**). In contrast, the C-terminal cytosolic tail exhibits relatively low sequence conservation (Kang et al., 2002) suggesting that cytosolic region is evolutionarily labile.

Identifiable orthologs to *cdon* and *boc* are present in non-vertebrate bilaterian taxa. These include *Drosophila* paralogs *ihog* and *boi*, whose function in hedgehog signaling has been well characterized (Bilioni et al., 2013; Camp et al., 2010; Yao et al., 2006). However, the orthologous relationships between *cdon*/*boc* and non-vertebrate orthologs such as *ihog* and *boi* are less well known. To explore the molecular evolution of this gene family we built a maximum likelihood tree (phyml) using 44 amino acid sequences from taxa representative for major bilaterian clades (**Figure 1B**). These data identify *cdon*/*boc* as shared vertebrate specific paralogs that emerge from a duplication event at either the base of the vertebrate phylogeny or along the gnathostome lineage following splitting from cyclostomes (hagfish and lamprey). Importantly, these data are consistent with a hypothesis that *cdon*/*boc* originated during one of the two vertebrate specific genome duplications (Dehal and Boore, 2005). Analyses suggest that an independent gene duplication event led to the origins of *boi* and *ihog* paralogs in drosophilids (**Figure 1B**)(Allen et al., 2011; Izzi et al., 2011; Sanchez-Arrones et al., 2012), a finding that is concordant with gene trees made publicly available by the TreeFam consortium (Li et al., 2006; Ruan et al., 2008).

To explore the function of *cdon*/*boc* we used mutations that target the conserved functional extracellular domains (**Figure 1C**, see methods). Two mutants for each gene were initially used to confirm an association between phenotypes and gene mutations. As we observed similar phenotypes for both *cdon* mutations and both *boc* mutations, we focused our analyses on two mutations in particular, cdon^co1019^ and boc^co25^, and refer to these mutants simply as *cdon* and *boc* except where stated.

### cdon and boc transcripts are expressed in tNCCs and slow-twitch muscle cells

To determine where endogenous *cdon* and *boc* transcripts are expressed we used multiplexed hybrid chain reaction *in situ* hybridization (HCR).

We found *cdon* and *boc* transcripts to be abundant in the dorsal central nervous system (CNS) of 24hpf-48hpf zebrafish embryos (**Figure 2A-D**), consistent with similar data from prior studies (Bergeron et al., 2011; Cardozo et al., 2014; Kearns et al., 2021). This pattern is established early in development where *cdon* and *boc* are expressed along the border of the neural plate in cells that contribute to the dorsal neural tube/NCC (**Figure 2A**). While *boc* transcripts in the CNS form a broad dorsal-ventral gradient (Kearns et al., 2021), *cdon* transcripts are restricted to the dorsal-most roofplate cells and the ventral-most floorplate cells of the CNS by 24hpf. In this way, *cdon* creates a dorsal sub-domain of *boc* expression (**Figure 2D**). Floorplate cells also express hedgehog ligand, and thus unlike *boc, cdon* is expressed in hedgehog ligand producing and non-producing cells. This CNS expression at 24hpf is similar to expression patterns reported in mouse embryos (Izzi et al., 2011; Kang et al., 2002; Mulieri et al., 2002; MULIERI et al., 2000; Tenzen et al., 2006).

**Figure 2.**
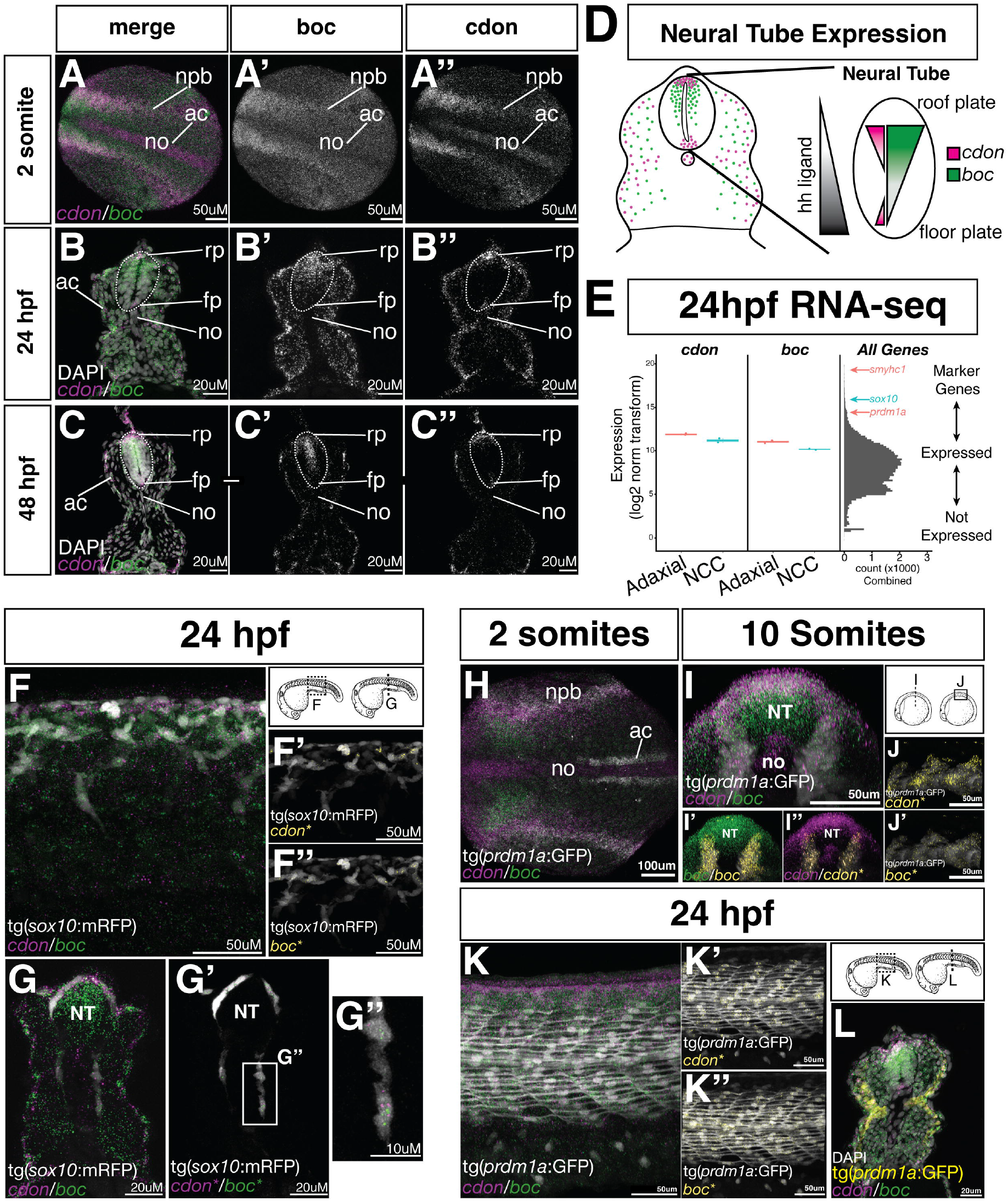
*cdon* and *boc* are expressed by NCCs and adaxial mesoderm cells. **(A-C)** HCR shows *cdon* (magenta) and boc (green) expression in the developing neural tube of 2 somite **(A)**, 24 hpf **(B)**, and 48 hpf **(C)** embryos. Panels **B&C** are 20µM sections at the level of the yolk extension (YE) in the trunk. Note expression at the border of the neural plate **(A)**, and in the rdorsal neural tube at 24 and 48 hpf. Weak expression is observed throughout the mesoderm. **(D)** Schematic summarizing expression of *cdon*/*boc* in the neural tube. Gradient of hedgehog ligand is indicated at left in grey, while expression of *cdon*/*boc* are indicated by magenta and green respectively. **(E)** Bulk RNA-seq from FAC-sorted tNCCs (blue) and slow-twitch muscle (red) suggest both tissues express *cdon*/*boc*. Boxplots show *cdon*/*boc* expression levels in 2 replicate samples. Right panel shows histogram of gene expression levels for all samples for all genes. Location of genes that mark NCCs or slow-twitch muscle are indicated. **(F&G)** HCR expression of *cdon*/*boc* in tNCCs. **(F)** whole mount lateral view of 24 hpf embryo showing expression of *cdon*/*boc* with NCCs (grey) labeled by the tg(*sox10*:mRFP) transgene. HCR puncta overlapping the *sox10* transgene (*) are segmented out and shown in yellow **(F’&F’’) (G)** Cross section shows tNCCs expressing *cdon*/*boc* transcripts, with HCR puncta overlapping the sox10 transgene **(G’ and G’’). (H-L)** Slow-twitch muscle cells express *cdon* and *boc*. The tg(*prdm1a*:eGFP) transgene labels adaxial cells at 2 somites **(H)**, and slow-twitch muscle cells at later stages **(I-L)**. (**I)** 3D rendered cross section of whole mount imaged embryo shows HCR puncta overlapping the tg(*prdm1a*:eGFP) transgene **(I’** and **I’’). (J)** 3D rendered lateral views show HCR puncta overlapping the *prdm1a* transgene. **(K)** 24hpf slow-twitch muscle cells (grey) continue to express *cdon*/*boc* transcripts. Panels **K’** and **K’’** show overlapping puncta. **(L)** Cross section of the trunk of a 24hpf embryo shows *cdon*/*boc* puncta in slow-twitch muscle.

Prior studies have suggested that *boc* is expressed by slow-twitch muscle mesoderm at 24hpf (Bergeron et al., 2011), however detailed investigation of *cdon/boc* expression in NCCs and slow-twitch muscle are not available. To further characterize expression, we used the tg(*sox10*:mRFP) and tg(*prdm1a*:eGFP) transgenic reporter lines to isolate both tissues from the trunks of 24hpf zebrafish embryos and sequenced the transcriptome of these tissues using bulk RNA-seq (see methods). These data identify *cdon*/*boc* transcripts in both tissues supporting the hypothesis that tNCCs and adaxial mesoderm express *cdon* and *boc* at low to moderate levels (**Figure 2E**).

We then asked if this expression is observable by HCR. Using the tg(*sox10*:mRFP) transgene to label NCCs, we segmented out *cdon* and *boc* HCR puncta that overlap the *sox10* transgene confirming *cdon*/*boc* transcripts in tNCCs at 24hpf (**Figure 2F&G**), albeit at levels far lower than the surrounding tissues.

To confirm expression in slow-twitch muscle/adaxial cell mesoderm, we used the tg(*prdm1a*:eGFP) (Elworthy et al., 2008). At the 2 somite stage, adaxial cells are labeled with the tg(*prdm1a*:eGFP) transgene, but we did not observe significant expression of *cdon*/*boc* in these cells (**Figure 2H**). By the 10 somite stage, *cdon* and *boc* transcripts were observed throughout the paraxial mesoderm. Using the tg(*prdm1a*:eGFP) transgene to segment out HCR transcripts expressed by adaxial cells, we show *cdon* and *boc* HCR puncta in slow-twitch muscle progenitor cells (**Figure 2I-J**). At 24hpf, differentiating slow-twitch muscle cells continue to express both *cdon* and *boc* (**Figure 2K-L**) consistent with findings from our bulk RNA-seq data (**Figure 2E**).

Taken together, these analyses indicate that *cdon* and *boc* are broadly expressed by mesodermal tissues that include adaxial cell derived slow-twitch muscle cells, as well as tNCCs at low levels, in addition to the developing neural tube.

### tNCC migration and slow-twitch muscle development are affected in cdon;boc mutants

To determine whether *cdon* and *boc* mutations affect tNCC migration and slow-twitch muscle development, we used the tg(*sox10*:mRFP) line to label migrating tNCCs and immunostaining of a slow-twitch muscle specific myosin (DSHB F59) to label adaxial mesoderm derivatives in 24hpf embryos.

In wildtype embryos, tNCCs collectively migrate in streams down the middle of each body segment in concert with the lateral migration of slow-twitch muscle cells (**Figure 3A**,**B**). We found that tNCC migration was normal in both single *cdon* and *boc* mutant embryos (**Figure 3D-E**). These data from homozygous mutant embryos are in contrast to those reported by morpholino knockdown of *cdon* (Powell et al., 2015). However, in double *cdon;boc* mutants, tNCCs failed to form distinguishable streams, and tNCC migration arrested at the ventral neural tube/body midline (**Figure 3F**), a phenotype similar to that observed in *cdon* morphant embryos and suggestive of genetic compensation between paralogs (Powell et al., 2015).

**Figure 3.**
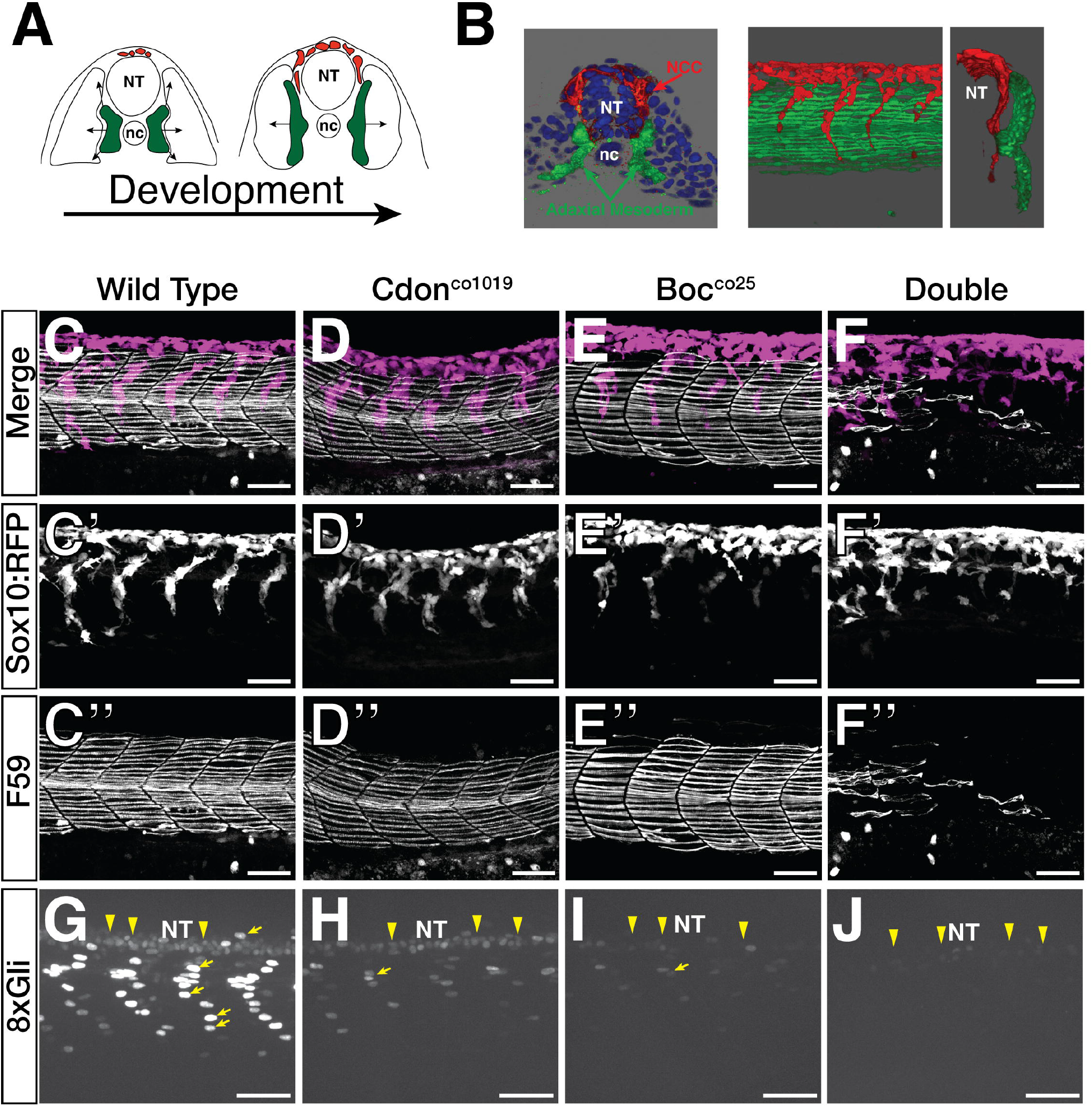
*cdon* and *boc* double mutants have NCC migration and slow-twitch muscle differentiation phenotypes and a reduction in hedgehog signal reception. **(A)** Schematic cross sections show NCC (red) and adaxial mesoderm (green) migration from approximately 10 somites to 20 somites. Adaxial mesoderm cells are specified adjacent to the notochord (nc) and migrate laterally to differentiate into slow-twitch muscle fibers. Concurrent with adaxial cell migration, NCCs migrate ventrally from dorsal neural tube (NT). **(B)** 3D renderings of a 10uM cross section from the trunk of a 15 somite (left) and a whole mount 24hpf embryo (right) showing association between migrating adaxial mesoderm (green) and NCCs (red). **(C-F)** Maximum projection images show NCC streams (tg(*sox10*:mRFP)) and immunostaining for slow-twitch muscle (F59) for 24hpf wildtype embryos. Note loss of both streams and slow-twitch muscle in the double mutant. All images taken over the yolk extension in the trunk. **(G-J)** Maximum projection images of the tg(8xGli:mCherry) transgene in 24hpf embryos. Note mCherry positive nuclei in the slow-twitch muscle mesoderm (yellow arrows) and ventral neural tube (yellow arrowheads). mCherry signal progressively decreases in *cdon;boc mutants* reflecting lower hedgehog signal reception.

Similarly, we found that slow-twitch muscle differentiation/migration was normal in *cdon* and *boc* (Bergeron et al., 2011) single mutants. Only in double mutants did we observe defects in slow-twitch muscle differentiation (**Figure 3C-F**). Loss of slow-twitch muscle cell differentiation was not complete, however, and posterior body segments tended to have fewer slow-twitch muscle fibers than anterior segments. In double mutant embryos, slow-twitch muscle fibers appeared dysmorphic with loosely packed myosin bundles relative to wildtype embryos (e.g. compare myosin bundle width and packing in **Figure 3**).

Thus defects in tNCC migration and slow-twitch muscle cell differentiation are observed in *cdon*;*boc* double mutants.

### cdon and boc affect hedgehog signal transduction in slow-twitch muscle cells

Since *cdon* and *boc* have previously been implicated in hedgehog signaling (Allen et al., 2011; Zhang et al., 2006), we hypothesized that defects in both tNCC migration and slow-twitch muscle cell differentiation were due to loss of hedgehog signaling that is only apparent in the double mutants. To test this, we measured hedgehog signal reception by using the transgenic, tg(8xGliBS:mCherry-NLS-Odc1) reporter line (hereafter 8xGli:mCherry) that drives nuclear localized mCherry under the control of 8 tandem Gli binding sites (Mich et al., 2014). The mCherry fluorophore contains an Odc1 destabilization domain the facilitates proteasomal degradation to promote rapid fluorophore turnover in expressing cells.

At 24hpf, we observed mCherry positive nuclei in ventral neural tube cells and in slow-twitch muscle fiber nuclei (**Figure 3G**)(Kearns et al., 2021; Mich et al., 2014). We never observed mCherry positive nuclei in tNCCs. In *cdon* and *boc* mutants, the number and signal intensity of nuclei that are 8xGli:mCherry positive was lower as compared to wildtype siblings (**Figure 3G-J**). This decrease in signal was true for both cells in the neural tube and slow-twitch muscle. Importantly, 8xGli:mCherry intensity varied across genotypes, with *boc* mutants displaying lower 8xGli:mCherry signal than *cdon* mutants, and double mutants exhibiting lower signal than all other genotypes. These data suggest that *cdon* and *boc* mutations have decreased hedgehog activity in slow-twitch muscle cells, which is consistent with a hypothesis that perturbations to hedgehog signaling is responsible for the slow-twitch muscle differentiation phenotype in *cdon* and *boc* mutant embryos (Barresi et al., 2000; Devoto et al., 1996).

Interestingly, the tg(8xGli:mCherry) transgene reported substantively lower hedgehog signal reception in single *cdon* and *boc* mutant embryos that exhibit normal slow-twitch muscle differentiation. These data suggest that slow-twitch muscle development is robust to moderate perturbations to hedgehog signal. If this hypothesis is correct, we reasoned that *cdon* and *boc* mutants should sensitize embryos to further pharmacological manipulation of smoothened signaling with cyclopamine. To test this, we in-crossed double heterozygous *cdon*;*boc* parents and treated embryos with a subthreshold dose of 20uM cyclopamine from 50% epiboly to 24hpf. We then quantified the number of slow-twitch muscle fibers (F59) at 24hpf as a readout.

While single mutants treated with ethanol as a control had a similar number of slow-twitch muscle fibers as their wildtype ethanol treated siblings, single mutants treated with 20µM cyclopamine had significantly fewer slow-twitch muscle fibers than their similarly cyclopamine treated wildtype siblings (**Figure 4**). This pattern was even more striking for the double mutant. While double mutants treated with ethanol have an average of approximately 5-6 slow-twitch muscle fibers per segment, double mutants lost all slow-twitch muscle fibers when treated with 20µM cyclopamine. We noted qualitative changes to slow-twitch muscle morphology in cyclopamine treated *cdon* and *boc* embryos. Mutants treated with 20µM cyclopamine often exhibited gaps or missing slow-twitch muscle fibers in body segments, and muscle fibers were more likely to have loosely packed myosin.

**Figure 4.**
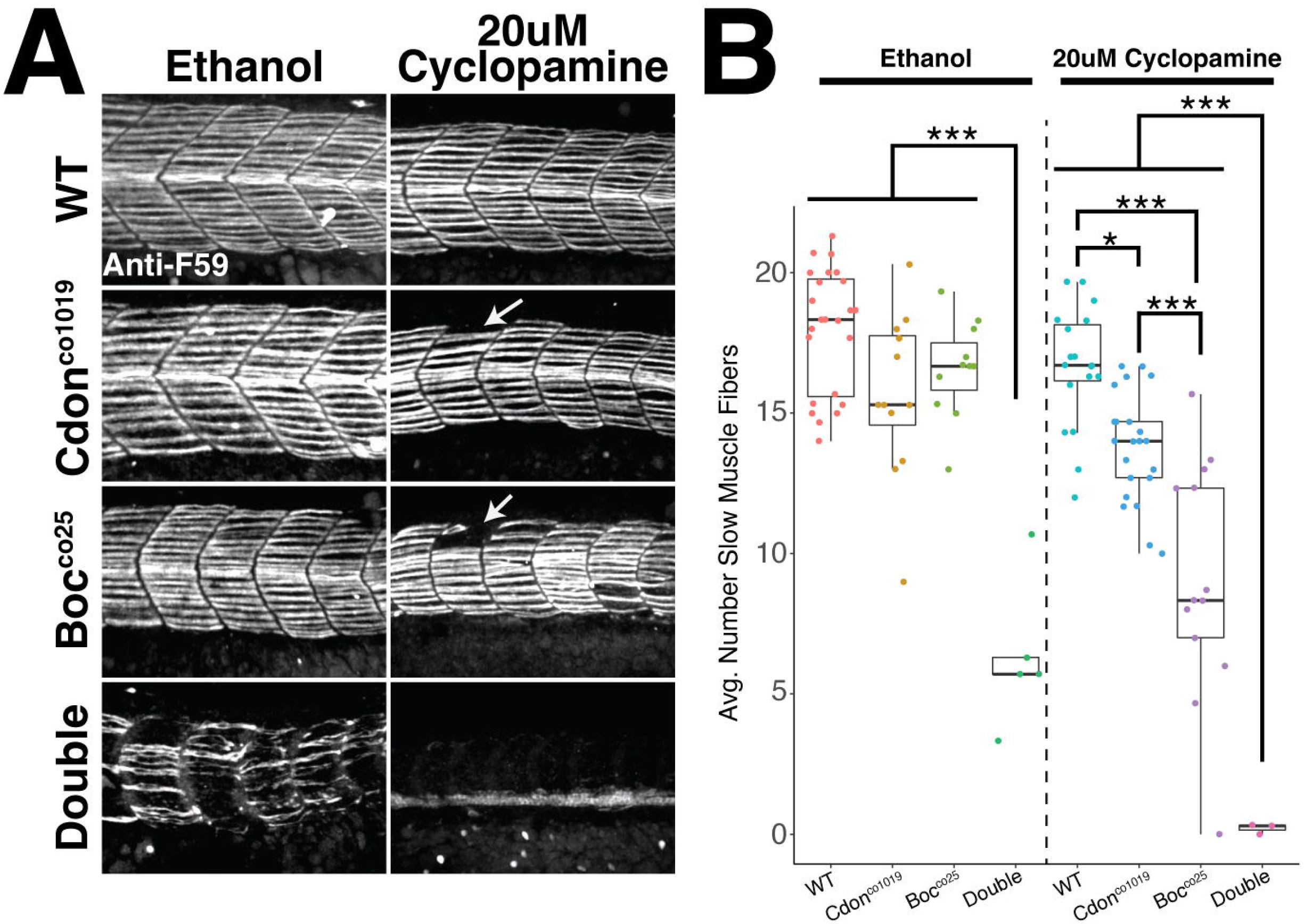
*cdon* and *boc* sensitize zebrafish embryos to a reduction in hedgehog signal by cyclopamine treatment. **(A)** Representative images show slow-twitch muscle fiber morphology in 24hpf embryos treated with either 20uM cyclopamine or an equivalent volume of ethanol as a control. Note *cdon* and *boc* mutants exhibit more severe phenotypes than their wildtype siblings when treated with cyclopamine as evidenced by gaps in segments (arrows), and complete loss of slow-twitch muscles in the double. **(B)** Quantification of average number of slow-twitch muscle fibers per segment shows that *cdon* and *boc* mutations are sensitizing embryos to cyclopamine treatment. Note that the number of slow-twitch muscle fibers in single *cdon* and *boc* mutants are similar to wildtype siblings when treated with ethanol, but significantly lower when treated with 20uM cyclopamine. Significance values are from Tukey pairwise post-hoc tests: <0.05 = *, <0.001 = ***.

We interpret these results to indicate that *cdon* and *boc* are modifying hedgehog signal reception by slow-twitch muscles, and that loss of *cdon* or *boc* sensitize embryos to further perturbations of hedgehog signal.

### Trunk neural crest cells lose directionality but not migratory ability in cdon;boc mutants

The arrested tNCC migratory phenotype in the double mutants is striking and could result from either a loss of NCC migratory ability or a loss of directionality. To investigate this further we used the tg(*sox10*:mRFP) transgene to label migrating tNCCs and live imaged NCC migration through the trunk of the zebrafish embryo beginning at 22hpf (**Figure 5**). Single tNCCs were then manually tracked. We found that across all genotypes, tNCCs were qualitatively migratory, and general measures of mobility (e.g. velocity etc.) in mutants were similar to that observed in wildtype controls (**Figure 5D-G**). In contrast, we observed a significant loss of directional NCC movement in double *cdon;boc* mutant embryos. In double mutants, tNCCs migrate to the body midline and then move along the anterior-posterior axis (**Figure 5A-C**), which is in contrast to the directed ventral movement typical of wildtype embryos. This is seen quantitatively as lower migratory directionality and an increased proportion of movements in lateral directions (**Figure 5B&H**). Single *cdon* or *boc* mutants did not display migratory phenotypes that were different from that of wildtype controls. Taken together, we interpret these results to indicate that loss of *cdon* and *boc* affects tNCC directionality and path-finding ability, but that *cdon* and *boc* are not required for overall tNCC mobility.

**Figure 5.**
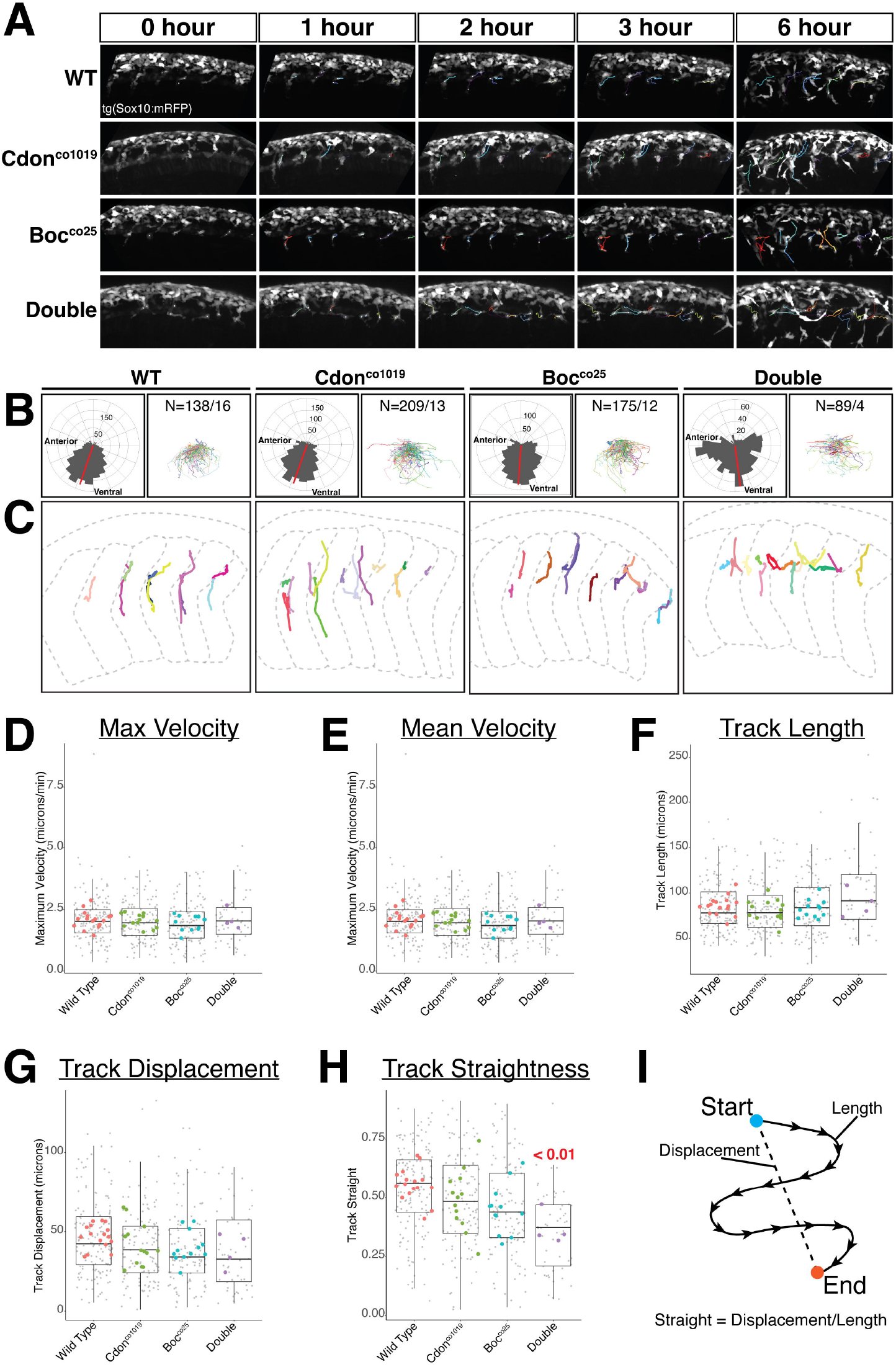
NCC directionality, but not migratory ability, is disrupted in the *cdon;boc* mutants. **(A)** Maximum projection images show migrating tNCCs in representative wildtype, *cdon, boc*, and double mutant embryos. Migratory tracks from individual cells are shown by colored lines. Note that NCCs form horizontal tracks at the body midline in the double mutant. **(B)** Matched roseplots (left panels) showing directional movement and line plots (right panel) showing tracks of NCCs in each genotype. For both panels, dorsal is up and anterior is to the left. Samples sizes reflect #tracks/#Embryos. Note that ventral directionality of NCC movement is lower in the double mutant as best seen by the rose plot. **(C)** NCC migration tracks (colored lines) are overlayed onto outlines of the body (grey dashed lines) for representative embryos. Note the vertical aligned tracks in the wildtype, *cdon*, and *boc* mutants that reflects ventral migration. Tracks form horizontal lines in the double mutant reflecting NCC migratory movements along the anterior-posterior axis. Body outlines do not reflect growth. Somite segments in the double mutant are wider and dysmorphic. **(D-H)** Boxplots show quantification of NCC migration for maximum velocity **(D)**, mean velocity **(E)**, track length **(F)**, track displacement **(G)**, and track straightness **(H)**. Note that the double *cdon;boc mutants* are the only condition that is significantly different from wildtype embryos in track straightness. Small grey points are individual track values, large colored points are means for individuals, boxplots reflect variability across tracks. **(I)** Schematic of track straightness which is calculated as track displacement dived by track length.

### cdon/boc affect tNCC migration non-cell autonomously

The observed defects in tNCC migration are similar to phenotypes observed in other hedgehog mutants that have previously been attributed to defects in slow-twitch muscle differentiation (Banerjee et al., 2011; Banerjee et al., 2013; Honjo and Eisen, 2005). We therefore hypothesized that tNCC migratory defects in *cdon*;*boc* mutants are a secondary consequence of failed slow-twitch muscle differentiation leading to defects in the secretion or deposition of tNCC guidance cues along the migratory path. In order to determine cell autonomous vs. non-cell autonomous function we attempted to rescue tNCC migration in double *cdon*;*boc* mutant embryos using two complementary methods.

First, we created genetic mosaics by transplanting double mutant NCCs into wildtype hosts (**Figure 6**) and examined tNCC migration. The rationale for these experiments is that if the observed tNCC migration defects are a non-cell autonomous consequence of slow-twitch muscle differentiation, then we would expect transplanted mutant tNCCs to migrate normally in wildtype hosts with normal muscle differentiation. This is exactly what we observed.

**Figure 6.**
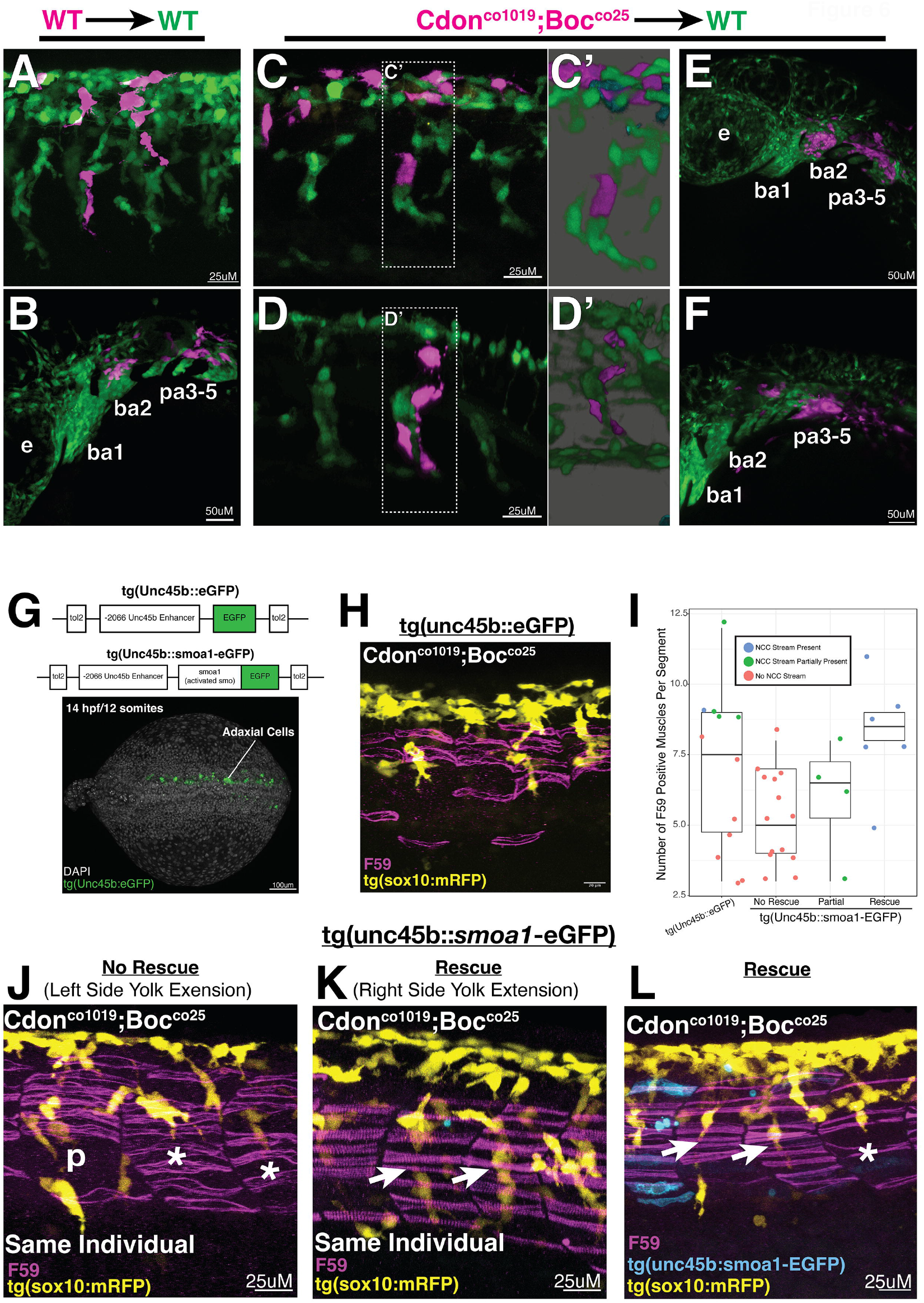
*cdon/boc* affect tNCC migration non-cell autonomously. **(A-F)** Mutant tNCCs transplanted into wildtype hosts migrate normally. For all transplants, donor embryos were derived by in crossing double heterozygous *cdon*;*boc* parents where 1/16 will be double mutants. All donor embryos were transgenic for the tg(*sox10*:mRFP) transgene, and wildtype embryos carry the tg(*sox10*:eGFP) transgene. Wildtype NCCs transplanted into a wildtype background migrate normally in both the trunk **(A)** and cranial **(B)** regions. Similarly, double mutant *cdon*;*boc* tNCCs migrate normally in wildtype hosts when transplanted into the trunk (C&D, 3/3 embryos) and cranial regions. For panels **(C**,**D)**, trunk streams are shown as both 3D projections and 3D volume renderings **(C’ & D’). (G-L)** Mosaic expression of *smoa1-eGFP* in adaxial cell progenitors leads to a partial rescue of tNCC streams. **(G)** tol2 constructs used for rescues. Dorsal view of a 10 somite embryo shows mosaic expression in adaxial cell progenitors. **(H)** Representative image of slow-twitch muscle morphology (F59) and tNCC streams in a double cdon/boc embryo injected with a control eGFP construct. **(I)** Number of slow-twitch muscle fibers per segment in eGFP/control and *smoa1a-eGFP* injected embryos. For *smoa1a-eGFP* injected embryos, points are separated by whether the tNCC stream in that segment was qualitatively rescued. IControl injected, N=4 embryos. *smoa1a-eGFP* injected, N=8. **(J-L)** Representative images of *smoa1a-eGFP* injected embryos. Panel **(J)** shows left side of an embryo where tNCC migration was not rescued, while panel **(K)** shows the right side of this same embryo where tNCC migration was rescued. Note the slow-twitch muscle fiber morphology is only rescued on the right side of the embryo. Panel **(L)** shows *smoa1a-eGFP* expression as mosaic. Arrows points to segments with qualitatively rescued tNCC streams. Asterisks label segments with non-rescued streams, “p” labels a representative “partial rescue”. e = eye, ba1 = branchial arch 1, ba2 = branchial arch 2, pa3-5 = posterior arches.

When wildtype cells from the region forming tNCCs are transplanted into wildtype hosts we observed transplanted tNCCs to migrate in well-organized streams alongside host tNCCs (**Figure 6A**). Similarly, in 3 of 3 cases where double mutant tNCCs were transplanted into wildtype hosts we observed mutant tNCCs migrating alongside host tNCCs in organized streams (**Figure 6C&D**). Cranial NCCs also migrated normally in all transplant conditions (**Figure 6B,E,F**). These data suggest that double *cdon/boc* mutant tNCCs migrate normally in wildtype hosts with normal mesoderm development suggesting a non-autonomous role for *cdon* and *boc* in tNCC migration.

The second approach used tol2 transgenesis to rescue the *cdon*/*boc* mutations in a tissue-specific manner. To do this, we used a -2.066kb fragment of the *unc45b* enhancer that drives expression in adaxial cell mesoderm at early segmentation stages (Berger and Currie, 2013). Importantly, this early adaxial cell expression turns on prior to slow-twitch muscle cell differentiation (**Figure 6G**). We also took advantage of a constitutively active smoothened (*smoa1*-eGFP) protein (Ju et al., 2014; Ju et al., 2009), reasoning that if *cdon*/*boc* are affecting hedgehog signaling then driving *smoa1*-eGFP in the slow-twitch muscle progenitor population should rescue both slow-twitch muscle morphology and tNCC migration (**Figure 6G**).

Out of 8 double mutant embryos injected with the *smoa1a*-egfp construct, 4 exhibited at least one tNCC stream that was qualitatively normal. This is in contrast to 4 double mutant embryos injected with a control eGFP construct where only 1 exhibited a tNCC stream that could be considered as a wildtype phenotype.

Importantly, these are F0 injected embryos, and thus tol2 transgenesis is mosaic, and the rescue of phenotype is also expected to be mosaic in the embryo. For instance, in one embryo injected with the *smoa1*-egfp construct, the left side of the embryos exhibited a typical mutant phenotype, while the right side of the embryo exhibited tNCC streams that appeared wildtype-like (**Figure 6J&K**). Further, the slow-twitch muscle morphology on the right side of this embryo was similarly wildtype-like, while the non-rescued left side appeared dysmorphic with loose packed myosin bundles.

To account for mosaic rescue, we analyzed 3 body segments over the yolk extension for each embryo by (1) qualitatively categorizing tNCC streams into rescue, partial rescue, or no-rescue categories based on tNCC stream formation, and (2) counting the number of slow-twitch muscle fibers (F59) for that segment. These data show that body segments with rescued tNCC streams tended to have more slow-twitch muscle fibers than either non-rescued or partial rescued segments (**Figure 6I**). Further, we noted that slow-twitch muscle fiber morphology appeared normal in these ‘rescued’ segments (note tight packed slow-twitch muscle fibers in **Figure 6** panels **K&L** compared to loosely packed fibers in panels **H&J**).

We did notice, however, that some control injected embryos had similar numbers of slow-twitch muscle fibers to those observed in the *smoa1*-eGFP injected embryos (**Figure 6I**, note range for control construct). Further, these body segments with greater numbers of slow-twitch muscle cells in the controls were more likely to exhibit ‘partial rescues’, or in one case a wildtype-like tNCC stream. We thus hypothesized that there is likely some degree of incomplete penetrance in the *cdon*;*boc* mutants whereby occasional segments in some embryos exhibit wildtype phenotypes. In this scenario, we interpret our mosaic expression of *smoa1*-eGFP to be pushing *cdon;boc* mutants towards a wildtype phenotype that is interacting with background variability (genetic or stochastic) in the penetrance of the *cdon*/*boc* alleles. Nonetheless, these data support the idea of a non-cell autonomous role for *cdon/boc* in regulating directed tNCC migration.

### Deposition of a Col1a1a extracellular matrix is affected in cdon;boc mutants

Prior research suggests that slow-twitch muscle cells express or modify extracellular matrix proteins including Col15a1b (Guillon et al., 2016), tenascin (Schweitzer et al., 2005), *col18a1* (Banerjee et al., 2013; Schneider and Granato, 2006), and collagen modifying enzyme *plod3*/*lh3* (Schneider and Granato, 2006). Unlike cranial NCCs, trunk NCCs are known to migrate on collagen substrates (Lallier et al., 1992) suggesting that changes to a collagen extracellular matrix (ECM) may be responsible for mediating the effects of *cdon*/*boc* on tNCC migration (Banerjee et al., 2013).

To investigate this further we mined our RNAseq dataset of FAC sorted slow-twitch muscle cells to find all collagen genes expressed by this tissue. These data identify many low to moderate collagen genes expressed by slow-twitch muscle cells. In particular, *col1a1a* is highly expressed by slow-twitch muscle, with more than twice the number of estimated transcripts than the next two most highly expressed collagen genes (**Figure 7A**).

**Figure 7.**
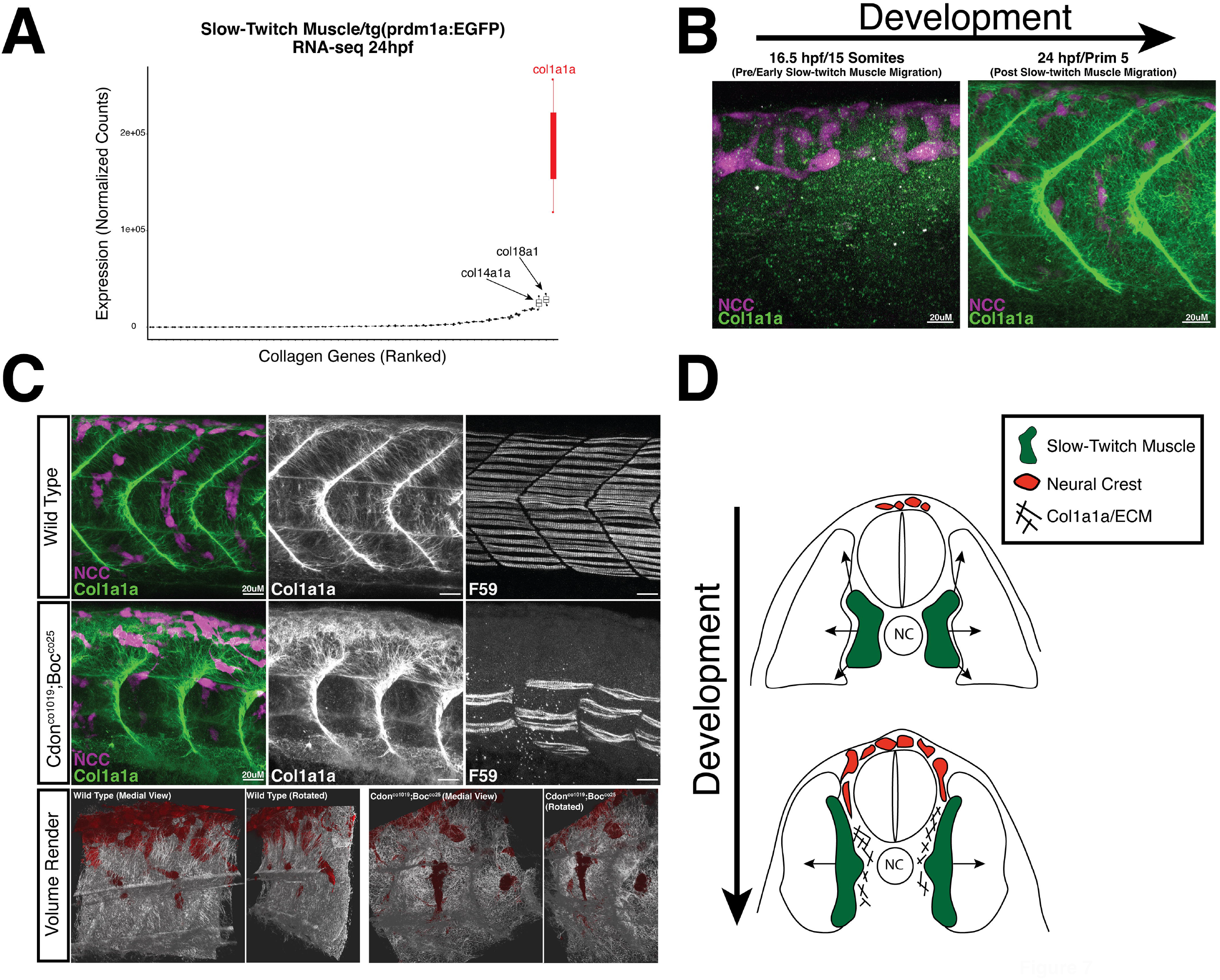
Col1a1a extracellular matrix (ECM) deposition is disrupted in cdon;boc mutants. **(A)** RNA-seq data from FAC sorted tg(*prdm1a*:eGFP) slow-twitch muscle cells at 24hpf identify numerous expressed collagen genes including *col1a1a*. **(B)** 3D projections of a 15somite and 24hpf embryo show Col1a1a protein deposition. **(C)** 3D projections (top rows) and volume renderings (bottom row) show medial Col1a1a deposition in wildtype and double mutant embryos. Note loss of Col1a1a ECM along the medial zone of each segment in the double mutant associated with loss of slow-twitch muscle differentiation. Retention of Col1a1a around dorsal neural tube (yellow arrows) coincides with location tNCCs often arrest migration. **(D)** Schematic model whereby slow-twitch muscle cells deposit an ECM containing Colo1a1a necessary for tNCC stream formation and migration.

We looked at Col1a1a protein deposition using an antibody raised against the zebrafish Col1a1a protein (GeneTex). In wildtype embryos at 24hpf, Col1a1a deposition was observed in the presumptive skin ectoderm, vertical myosepta, and medial border of the mesoderm adjacent to the neural tube and developing gut (**Figure 7B**). At 16.5 hours, we did not observe Col1a1a fibers, suggesting that this ECM component is deposited after this time period (Figure 7B). Intriguingly, this period between 16.5hpf and 24hpf coincides with slow-twitch muscle migration (Elworthy et al., 2008).

Importantly, we observed Col1a1a fibers deposited as ECM along the medial path that tNCCs migrate on (**Figure 7C**). In wildtype embryos, Col1a1a is richly deposited along the vertical myosepta, and fibers extend into the medial region where tNCC streams are observed. However, in double *cdon*;*boc* mutants, Col1a1a deposition in the tNCC stream pathway was largely absent. Col1a1a was present in other areas of the embryos such as the vertical myosepta and skin suggesting that loss of Col1a1a in the trunk is specific to the tNCC stream path region (**Figure 7C**).

We interpret these data to suggest that tNCC migratory defects may be a consequence of loss of the Col1a1a ECM along the migratory path (**Figure 7**). Supporting this hypothesis is the observation that Col1a1a is still expressed dorsally in the double mutants, but stops at the position where tNCCs often arrest. Thus the arresting migratory phenotype and anterior-posterior movements of tNCCs along this axis are hypothesized as a consequence of the loss of Col1a1a expression in the ventral-medial compartments of each body segment.

## Discussion

### cdon and boc affect tNCC migration non-cell autonomously

Here we show that *cdon* and *boc* affect tNCC migration and differentiation of slow-twitch muscle cells. Despite our prior knockdown studies suggesting that *cdon* alone functions in tNCC migration (Powell et al., 2015), we find that only in double *cdon;boc mutants* do we observe noticeable tNCC migratory defects that are associated with loss of slow-twitch muscle. Prior research suggests that slow-twitch muscle differentiation requires hedgehog signaling (Elworthy et al., 2008), and that tNCC migration requires proper slow-twitch muscle differentiation (Honjo and Eisen, 2005). Our data are consistent with these result and suggests that the tNCC migratory defect in *cdon;boc* mutants is a consequence of defects in mesodermal differentiation (Banerjee et al., 2011; Banerjee et al., 2013; Honjo and Eisen, 2005). This hypothesis is further supported by the rescue experiments, that emphasize the importance of non-autonomous adaxial cell mesoderm differentiation for proper neural crest migration in zebrafish.

While mesodermal tissues are appreciated as critical for neural crest migration (Krull, 2010; Kulesa and Gammill, 2010; Theveneau and Mayor, 2012), the mechanisms underlying how the mesoderm affects NCC migration vary among both taxa and different NCC subpopulations. In many tetrapod taxa, mesoderm restricts cranial and trunk NCC migration along conserved paths through neuropilin, semaphorin, and ephrin ligand-receptor interactions (Davy and Soriano, 2007; Gammill et al., 2007; Gammill et al., 2006; Krull, 2010; Krull et al., 1997; Santiago and Erickson, 2002). Thus, the localization of receptors and ligands within tissues creates permissive/non-permissive zones that restrict NCC migratory paths. In addition, mesodermal tissues can act as a source for chemoattractant molecules such as chemokine signaling (Kasemeier-Kulesa et al., 2010; Olesnicky Killian et al., 2009). In zebrafish, semaphorin/neuropilin interactions mediate cranial NCC migration (Yu and Moens, 2005), though the role of this pathway in zebrafish tNCC migration is less established. To identify complementary ligand-receptor pairs in FAC sorted tNCCs and slow-twitch muscle cells, we searched our RNA-seq data. While some possible interacting molecules could be found, there were no clear candidate signaling pathways, leading us to hypothesize that there is not a direct ligand-receptor interaction between tNCCs and slow-twitch muscle mesoderm (not shown, accessible in NCBI Geo:GSE213728).

Instead, we build on prior research in both tNCC migration (Banerjee et al., 2011; Banerjee et al., 2013) and motor axon guidance (Guillon et al., 2016; Schneider and Granato, 2006; Zeller and Granato, 1999; Zhang et al., 2004) to suggest that slow-twitch muscle cells affect tNCC stream formation through ECM deposition. Adaxial derived slow-twitch muscle cells express Col15a1b (Guillon et al., 2016), *col18a1* (Schneider and Granato, 2006), as well as chondroitin sulfate proteoglycans associated with collagen-rich ECMs (Zhang et al., 2004). Morpholino knockdown of *col18a1* leads to tNCC migratory defects that are similar to those we observe in *cdon;boc mutants* (Banerjee et al., 2013). Our data add to this literature, showing that slow-twitch muscle express other collagen genes in addition to those previously studied, including *col1a1a* at exceptionally high levels. Deposition of *col1a1a* along the medial somitic boundary is lost in *cdon;boc mutants* consistent with a model where slow-twitch muscle cells are responsible for depositing a collagen rich ECM along the tNCC migratory path.

How the ECM regulates NCC migration is an active area of research (Leonard and Taneyhill, 2020; Vega-Lopez et al., 2017). One hypothesis is that ECM components create a permissive zone that bias NCCs to move on specific migratory routes (Banerjee et al., 2013; Leonard and Taneyhill, 2020). Our hypothesis of a collagen ECM creating a permissive migratory route is supported by chick neural tube explant studies, which show that trunk, but not cranial NCCs utilize collagen ECMs as preferred migratory substrates (Lallier et al., 1992). Thus slow-twitch muscle cells may be depositing a complex collagen-rich ECM critical for tNCC stream formation. However, other mechanisms may also be at play. For instance, cranial NCCs utilize durotaxis to migrate along a stiffness gradient of pre-chordal mesoderm in *Xenopus* (Shellard and Mayor, 2021). Muscle segment organization is disrupted in different slow-twitch muscle mutants (Henry and Amacher, 2004; Snow and Henry, 2009) and thus mesoderm stiffness may be affected in the *cdon;boc mutants* that exhibit defects in slow-twitch muscle differentiation. Whether tNCCs utilize durotaxis has not been studied to our knowledge.

### A possible cell autonomous hypothesis for cdon/boc function

Though we argue that *cdon*/*boc* affect tNCC migration non-cell autonomously through slow-twitch muscle differentiation, *cdon* and *boc* may also play cell autonomous roles in NCC migration. Indeed, NCC autonomous and non-autonomous function are not mutually exclusive hypotheses. We point to data showing *cdon/boc* expression in both the neural plate border, migrating tNCCs and mesoderm to suggest multiple roles for these genes. Recent work in zebrafish showed that notch signaling among tNCC pre-migratory progenitors establishes a unique ‘leader cell’ identity that is required for stream formation (Alhashem et al., 2022; Richardson et al., 2016) (McLennan et al., 2015; Morrison et al., 2017). Intriguingly, notch mutant embryos that lack leader cell identity have normal slow-twitch muscle differentiation and tNCC migratory phenotypes that are remarkably similar to what we observe in *cdon;boc mutants* (Alhashem et al., 2022). Whether *cdon*/*boc* also play a role in establishing a ‘leader cell’ identity, or function in ‘leader cell’ guidance is unknown. It is worth noting that our transplant rescue experiments cannot differentiate between a role for mesoderm or ‘leader cell’ identity, as leader cells will be present in the wildtype hosts.

### cdon and boc as modifiers of hedgehog signaling

Our data add to a growing literature showing that *cdon* and *boc* act synergistically during zebrafish development to modify hedgehog signal reception and regulate the development of multiple traits (Echevarria-Andino and Allen, 2020; Song et al., 2015; Zhang et al., 2011). Interestingly, we found that *cdon* single mutants do not have noticeable phenotypes in zebrafish, and we can recover adult homozygous mutant zebrafish (not shown). This is contrast to mouse, where loss of CDON results in mild holoprosencephaly (Allen et al., 2011; Allen et al., 2007; Echevarria-Andino and Allen, 2020; Zhang et al., 2011). While it is possible that our *cdon* mutations do not result in full loss of function, we find this possibility unlikely. Both *cdon* mutations lead to a predicted early stop prior to the transmembrane domain, we cannot fully rule out the possibility that a late start rescues *cdon* protein resulting in a hypomorphic but functional receptor or creates a dominant negative. Future research could investigate these possibilities.

An alternative possibility is that *cdon* and *boc* interact with other hedgehog pathway members resulting in species specific phenotypic responses. Thus in zebrafish other genes may provide robustness against *cdon* mutations and act as a buffer. The cyclopamine pharmacological experiments support this hypothesis. Intriguingly, *cdon* expression is often nested within *boc* expression suggesting that *cdon* and *boc* act together to modify hedgehog morphogen gradients. Kearns et al. 2021 found that *patched-2* (*ptch2*) expression in the dorsal neural tube is expanded in *boc*^*co25*^ mutant embryos, suggesting that loss of *boc* alone leads to more dorsal hedgehog reception (*ptch2* is a readout for hedgehog activity). These data were interpreted to suggest that *boc* sequesters hedgehog ligand driving hedgehog signal reception to ventral tissues. Expanding on this hypothesis, we note that *cdon* transcripts are positioned even more distally in the neural tube from hedgehog ligand source than are *boc* transcripts. Thus together *boc* and *cdon* gradients may act as strong sequestering agents to limit hedgehog signal activation and inhibit ligand activity in dorsal neural tube/NCCs.

<u>In summary,</u> we show that *cdon* and *boc* act synergistically in zebrafish to regulate slow-twitch muscle cell differentiation and affect tNCC migration non-cell autonomously through regulation of ECM deposition. Our data add to a growing literature in zebrafish showing the critical role of the slow-twitch muscle for regulating migration of cells. Intriguingly, homologous adaxial/slow-twitch muscle tissues in other taxa are less understood (Grimaldi et al., 2004) (Hammond et al., 2009), and to our knowledge there is no known tissue in amniotes that is homologous to adaxial cells. Despite this, *cdon* and *boc* are known to be critical for myogenesis in amniotes (Jeong et al., 2017; Lu and Krauss, 2010), and genes expressed by the slow-twitch muscle in zebrafish also regulate tNCC migration in mouse (Banerjee et al., 2011). Further, we and others note that *cdon* and *boc* are expressed in a broad range of tissues including the mesoderm and NCCs among others. *cdon*, but not *boc*, transcripts are also co-expressed in hedgehog ligand producing cells, and some studies have suggested that *cdon* plays a role in presenting hedgehog ligand to responsive cells (Hall et al., 2021). Thus it is likely that *cdon* and *boc* are playing different roles in different tissues, and that cellular context, expression levels, and distance from ligand sources, may be critical for determining the biological function of these receptors.

## Materials and Methods

### Fish stocks, husbandry, and generation of mutants

Adult fish and embryos were maintained at the University of Colorado Anschutz Medical school according to standard zebrafish rearing protocols and guidelines for the rearing and maintenance of vertebrate animals. Two *boc* mutations were investigated. The *boc*^co25^ mutant was previously recovered from an ENU screen (Kearns et al., 2021) and results in an non-synonymous isoleucine to serine substitution in the third FNIII domain (**Figure 1C**). The boc^uml/ty54z^ mutant was acquired from the Zebrafish International Resource Center and results in a predicted early stop (Bergeron et al., 2011). Both mutations are previously characterized as loss of function.

We used CRISPR/Cas9 to generate novel mutations in *cdon*. Two sets of guide RNAs (sgRNA) were designed to target either the 2nd and 3rd exons (ggcagUUgaggagcagagUa, UggUaggagccUgUgagagc, UgcUUcagaaUcgcUgagag), or to target the 11^th^ exon (UcgccaUUgagaUUgUgggg, cUccacgcgaaaUgcagUga) of cdon. DNA oligos were annealed to a common backbone scaffold, which was used to generate sgRNAs for injections (Williams et al., 2018). For each sgRNA set, our final CRISPR injection solution consisted of sgRNAs diluted to 20ng/uL, 1ug Cas9 Protein (Invitrogen TrueCut V2), and 5ug Rhodamine Dextran in a total volume of 10uL. Approximated 2-4nl were injected into the cell of a 1cell stage embryo, and F0 injected embryos were reared to adulthood.

We recovered a single allele for each sgRNA set. These include the cdon^co1019^ allele, which is a 229bp deletion that spans the second to third exon of *cdon*, removes the intronic sequence between these exons, and is predicted to result in an early stop (**Figure 1C**). The cdon^co1023^ allele is a 5bp deletion resulting in a predicted early stop in the 11^th^ exon that codes for the second FNIII domain. F0 injected embryos were outcrossed for a minimum of two generations to minimize effects of off target cutting.

Zebrafish *boc* and *cdon* mutants were crossed into existing transgenic reporter lines including the tg(−7.2kb*sox10*:mRFP)^vu234^ that labels NCCs with cytosolic mRFP (Kucenas et al., 2008), and the gli reporter tg(8xgli-Xla.Cryaa:NLS-d1mCherry) line that has 8 gli-binding sites driving a destabilized nuclear localized mCherry (Mich et al., 2014). The tgPAC(*prdm1a*:eGFP) line was used to label adaxial mesoderm for reference (Elworthy et al., 2008).

### cdon/boc molecular conservation and gene tree

Amino acid coding sequences for select bilaterian taxa were downloaded from NCBI GenBank. Orthologous sequences were searched directly or identified through BLAST searches and manual curation. The final set of sequences was chosen to include taxonomic representatives for major bilaterian taxa for which high confidence sequences are available. Effort was made to identify two paralogs for each taxon. Amino acid sequences were aligned using MUSCLE (v3.8.31) with default parameters (Edgar, 2004). Amino acid conservation (Figure 1A) was calculated by pairwise comparisons of *cdon*/*boc* homologues from 11 taxa (human, mouse, chicken, *Anolis*, frog, bichir, Gar, sheepshead minnow, medaka, catfish, zebrafish) and plotted in R by smoothing conservation values across 20 amino acid windows. Substitution models were selected using ProtTest3 by AIC (Darriba et al., 2011). A maximum likelihood gene tree was built using phyml (v3.3) and a JTT+I+G+F model with 1000 bootstraps (Guindon and Gascuel, 2003). Nodes with less than 60% bootstrap support were collapsed into polytomies. We recovered similar results from filtered alignments that removed poorly aligned regions and regions with gaps.

### Immunohistochemistry, in situ hybridization, and stains

Adaxial mesoderm derivatives were stained by fluorescent immuno-histochemistry against a slow-twitch muscle specific myosin (anti-MYH1A, F59, DSHB) at a concentration of 1:20 as previously described (Elworthy et al., 2008; Nguyen-Chi et al., 2012). Hybrid Chain Reaction (HCR) *in situ* hybridizations were performed as previously described (Choi et al., 2018; Lencer et al., 2021). Unlike traditional enzymatic methods for *in situ* hybridization, HCR allows single molecule measurement of gene expression at cellular resolution, which enables identifying cells and cell types that express transcripts at relatively low levels (Kearns et al., 2021). Custom HCR probes for zebrafish *cdon* and *boc* were acquired from Molecular Instruments and used at a concentration of 2picoMol/500uL Hybridization Buffer.

### RNA-seq of tNCCs and adaxial cell mesoderm

We used bulk RNA-seq to characterize the transcriptome of tNCCs and adaxial cell mesoderm at 24 hpf. To do this we took advantage of embryos that were doubly transgenic for the tg(*sox10*:mRFP) transgene that labels NCCs and tg(*prdm1a*:eGFP) transgene that labels adaxial mesoderm. Trunks from 80 embryos were dissected and cells were dissociated by incubating in Accumax (Stem cell technologies) at 37°C for 30 minutes with intermittent agitation. Cells were cleaned of debris by passing through a 40µM mesh filter, and washed in Dulbecco’s phosphate buffered saline plus DNaseI (final 7500U/mL). NCCs and adaxial mesoderm cells were isolated by FACS using a MoFlo XDP100 at the CU Anschutz Flow Cytometry Core.

Total RNA was extracted from FAC sorted samples of tNCCs and adaxial mesoderm using the RNAqueous-Micro Isolation Kit (ThermoFisher). Library preparation and Illumina sequencing (PE 150bp) was performed by the CU Anschutz Genomics core on a NovaSeq6000. Reads were aligned to the zebrafish genome (GRCz11, ensemble annotations) using STAR aligner (Dobin et al., 2013) and gene counts were calculated using featurecounts (Liao et al., 2014) with default parameters. DEseq2 (Love et al., 2014) was used to estimate normalized expression levels using the normTransform function.

### Imaging and tracking of migrating cells

For live-imaging and tracking, neural crest cells were labelled using the tg(*sox10*:mRFP) line. Embryos were mounted in 0.4% low melt agarose and neural crest cells were imaged over the yolk extension every 5 minutes from 22hpf to 30hpf on an Andor spinning disk confocal microscope at 28°C. Neural crest cells were manually tracked along the xy axis from maximum projection images using the ImageJ mTrackJ plugin (Meijering et al., 2012). Drift was accounted for using the Correct_3D_Drift.py plugin as implemented in ImageJ (Parslow et al., 2014). As we only observed a tNCC migratory defect in the medial migrating streams, we limited our tracking of cells to tNCCs migrating along this route. Quantitative analysis of track mobility was performed in R. A small number of short tracks with fewer than 30 points (2.5hrs) were excluded. As we also noted that NCCs often stopped migrating upon reaching a ventral position we further limited tracks to a maximum of 6 hours in order to capture cell migration. Limiting all tracks to the same time (e.g. number of points) produces similar results. Rose plots depict cell movements with a velocity greater than 0.3µM/min, which was empirically determined as a cutoff for capturing cell movements while excluding noise from drift and changes in cell shape. Using different cutoffs produced similar results. Linear models were built in R by including track length as a fixed effect, and embryo identity and date of imaging as random effects, with the packages lme4 (Bates et al., 2015), lmerTest (Kuznetsova et al., 2017), and emmeans. Images of tracks were produced in napari (Sofroniew et al., 2019).

### Transplant Rescue Experiments

Transplant rescues were performed by transplanting cells from gastrula stage embryos produced by in crossing double heterozygous *cdon*^*co1019*^*;boc*^*co25*^ parents into gastrula stage wildtype hosts. All donor embryos were transgenic for the tg(*sox10*:mRFP) transgene, while hosts were transgenic for the tg(*sox10*:eGFP) transgene (Carney et al., 2006), allowing us to differentiate transplanted NCCs (red) from host NCCs (green). Embryos were fixed at 24hpf, genotyped, and imaged for tNCC migration.

### Tol2 Rescue Experiments

Tol2 rescue experiments were performed following standard gateway cloning and tol2 transgenic methods (Kwan et al., 2007). Briefly,the primers 5’-GCGTAAAACTGTGGCGCGTAAAAC-3’ and 5’-CATTGAAGTCAATTCAGCTTCGTC-3’ were used to clone the Unc45b promoter from wildtype genomic DNA following (Berger and Currie, 2013). This produced a 2.066kb fragment of the 5’ upstream region to the *unc45b* gene. This fragment was placed into a tol2kit p5Entry vector using TA cloning (Invitrogen). A *smoa1*-eGFP construct was obtained as generous gift from Michael Taylor and subcloned into a tol2 middle entry vector. Expression constructs using the -2.066Unc45b enhancer to drive either *smoa1*-eGFP, or eGFP as a control, in adaxial cell progenitors.

For rescues, double heterozygous *cdon*^*co1019*^*;boc*^*co25*^ parents were in crossed and embryos were injected at the 1 cell stage with tol2 constructs and tol2 mRNA at a total injection volume of ∼2nL. Embryos were reared to 24hpf and then stained for F59 by immunofluorescence (see above) to assess tNCC stream formation and slow-twitch muscle cell morphology.. Wildtype and double mutant embryos were imaged over the yolk extension. Sick embryos were not used for analyses.

### Pharmacological manipulation

For pharmacological manipulation, embryos were manually dechorionated and treated with 20uM cyclopamine in ethanol (5uL/mL) from 50% epiboly to 24 hpf. Embryos that appeared developmentally delayed were excluded.

### Imaging, Statistical Analysis, and biological replication

All imaging was performed on either a Leica Sp8 LSM or Andor spinning disk confocal microscope. Image analysis and presentation was performed in ImageJ or napari. All statistical analyses were performed in R. Experiments were repeated at least 3 times and biological variation was accounted for in models and through inclusion of biological replicates.

## Acknowledgements

We thank members of the Artinger and Prekeris laboratories for comments on this manuscript and project. Special thanks to Caleb Doll, Bruce Appel, Christina Kearns for providing tools and advice. We thank Michael Taylor for his generous gift of the smoa1-eGFP construct. This work was in part supported by GM122768 (to RP) and a University of Colorado RNA Biosciences Initiative award to EL and KA. EL was supported by National Institutes of Health Ruth L. Kirschstein National Research Service Awards (T32CA17468 and F32HD103406). EL, RP, and KA conceived of the study and wrote the manuscript. EL performed research and analyses.

